# Accumulation of m^6^A exhibits stronger correlation with MAPT than β-amyloid pathology in an APP^NL-G-F^/MAPT^P301S^ mouse model of Alzheimer’s disease

**DOI:** 10.1101/2023.03.28.534515

**Authors:** Lulu Jiang, Rebecca Roberts, Melissa Wong, Lushuang Zhang, Chelsea Joy Webber, Alper Kilci, Matthew Jenkins, Guangxin Sun, Sherif Rashad, Jingjing Sun, Peter C Dedon, Sarah Anne Daley, Weiming Xia, Alejandro Rondón Ortiz, Luke Dorrian, Takashi Saito, Takaomi C. Saido, Benjamin Wolozin

**Affiliations:** Department of Pharmacology, Physiology and Biophysics, Chobanian and Avedesian School of Medicine, Boston University, Boston, MA, 02118; Department of Biological Engineering, Massachusetts Institute of Technology, Cambridge, MA 02139, USA; Department of Neurosurgical Engineering and Translational Neuroscience, Graduate School of Biomedical Engineering, Tohoku University, Sendai, Japan; Singapore-MIT Alliance for Research and Technology, Antimicrobial Resistance IRG, Campus for Research Excellence and Technological Enterprise, Singapore 138602, Singapore; Geriatric Research Education and Clinical Center, Bedford VA Healthcare System, Bedford, MA, 01730, USA; Laboratory for Proteolytic Neuroscience, RIKEN Center for Brain Science, Wakoshi, Saitama, 351-0198, Japan; Department of Neurology, Chobanian and Avedesian School of Medicine, Boston University, Boston, MA, USA; Center for Systems Neuroscience, Boston University, Boston, MA USA

**Keywords:** Tauopathy, Neurodegeneration, RNA binding proteins, RNA metabolism, TIA1, tau oligomers, tau fibrils, stress granules, neuropathology, RNA methylation, neuritic plaques.

## Abstract

The study for the pathophysiology study of Alzheimer’s disease (AD) has been hampered by lack animal models that recapitulate the major AD pathologies, including extracellular β-amyloid (Aβ) deposition, intracellular aggregation of microtubule associated protein tau (MAPT), inflammation and neurodegeneration. We now report on a double transgenic APP^NL-G-F^ MAPT^P301S^ mouse that at 6 months of age exhibits robust Aβ plaque accumulation, intense MAPT pathology, strong inflammation and extensive neurodegeneration. The presence of Aβ pathology potentiated the other major pathologies, including MAPT pathology, inflammation and neurodegeneration. However, MAPT pathology neither changed levels of amyloid precursor protein nor potentiated Aβ accumulation. The APP^NL-G-F^/MAPT^P301S^ mouse model also showed strong accumulation of N^6^-methyladenosine (m^6^A), which was recently shown to be elevated in the AD brain. M6A primarily accumulated in neuronal soma, but also co-localized with a subset of astrocytes and microglia. The accumulation of m6A corresponded with increases in METTL3 and decreases in ALKBH5, which are enzymes that add or remove m^6^A from mRNA, respectively. Thus, the APP^NL-G-F^/MAPT^P301S^ mouse recapitulates many features of AD pathology beginning at 6 months of aging.

## Introduction

The hallmark pathologies of Alzheimer’s disease (AD) consist of the accumulation of neuritic plaques composed of β-amyloid (Aβ), the accumulation of neurofibrillary tangles (NFTs) composed of microtubule associated protein tau (MAPT, Tau), inflammation and neurodegeneration [1]. Mutations in amyloid precursor protein (APP) cause AD, however cognitive loss is only weakly correlated with the accumulation of Aβ [1]. Cognitive loss is much more robustly correlated with MAPT based NFTs pathology, and mutations in MAPT are sufficient to cause dementia (frontotemporal dementia, FTD) [1]. Mutations in MAPT might not cause AD because MAPT pathology does not drive the accumulation of Aβ pathology [1]. The requirement of Aβ and MAPT pathologies to model AD has posed a challenge for mouse models of AD because mutations in either gene alone are insufficient to produce both pathologies in mice.

Models relying only on genetic modification of amyloid precursor protein (APP) develop abundant Aβ plaques but produce little MAPT pathology beyond modest increases in phosphorylation [2]. Mouse models over-expressing mutant APP, such as Tg2576, or over-expressing mutant APP and mutant presenilin 1, such as 5xFAD, rapidly develop accumulated Aβ and develop neuritic plaques. However, over-expressing APP and presenilin 1 cause effect resulting from the over-expression that are unrelated to the disease process [3, 4]. These problems have been addressed by knocking in the human APP gene containing mutations that increase production of Aβ40 and/or Aβ42 [3, 4]. These mice develop robust plaque pathology beginning as early as 3 months of age, however they develop little tau pathology and little neurodegeneration [3, 4].

The absence of robust MAPT pathology in mouse models expressing only endogenous MAPT likely derives from the low aggregation propensity of murine MAPT [5, 6]. This limitation has been addressed by introducing human tau constructs into the mouse brain [2, 7]. Many mouse models have been developed based on over-expressing wild type (WT) tau or mutant forms of tau linked to frontotemporal dementia [2, 7, 8]. These models develop robust tau pathologies, including pathologically phosphorylated, misfolded, oligomeric and/or fibrillar forms of tau pathology. These models also exhibit progressive neurodegeneration, which is consistent with observations in humans that cognitive loss is more closely associated with tau pathology than Aβ pathology [9-11]. Human MAPT knock-in models have also been developed, and these models produce tau pathology, but only very late in the murine life span (∼15 months) [2-4].

Multiple groups have explored crossing transgenic APP mouse over-expression models with tau mouse models [12-19]. The results generally show that Aβ accelerates tau pathology [12-14, 16]. The tau pathology does not appear to increase Aβ deposition [12-17]; indeed, in some cases it accelerates removal of Aβ [13]. These studies all suffer because they exhibit artifacts arising from the over-expression APP (WT or mutant) and in some cases mutant presenilins [3, 8, 20]. Recent studies have begun to explore crossing human knockin models [2]. Human APP knockin (KI) models develop robust Aβ pathology, yet do not exhibit artifacts associated with APP over-expression, such as elevated levels of APP cleavage products [3]. Human MAPT KI models exhibit delayed tau pathology, as does a cross between the two KI models [21, 22]. Thus, we sought to create a model that avoided artifacts associated with APP over-expression, yet still exhibited robust tau pathology.

We now report creating a mouse model in which the APP^NL-G-F^ KI and the PS19 P301S MAPT mouse lines were crossed. The resulting mouse model, termed APP^NL-G-F^/MAPT^P301S^, develops many aspects of AD pathology. The APP^NL-G-F^/MAPT^P301S^ mouse exhibits a progressive increase in Aβ load, neuritic plaques, all major forms of tau pathology, as well as exhibiting enhanced microglial activation, astrogliosis, neurodegeneration and a progressive loss in cognitive function. This model also exhibits other key elements of AD pathology including inflammation and elevated levels of N6-methyl-adenosine (m^6^A) tagged RNA, which has recently been shown to change strongly with disease progression [23]. In this APP^NL-G-F^/MAPT^P301S^ model, we also found that Aβ pathology potentiates MAPT pathology, astrogliosis, inflammation and neurodegeneration. MAPT pathology does not appear enhance the accumulation of Aβ pathology or inflammation beyond that observed in the APP^NL-G-F^ mouse. Intriguingly we found that levels of m6A RNA correlate with MAPT but not Aβ pathology.

## Methods

### Mice

Use of all animals was approved by the Boston University Institutional and Animal Care and Use Committee (IACUC). All animals used in this study were handled according to IACUC approved protocols and housed in IACUC approved vivariums at the Boston University Animal Science Center. The APP^NL-G-F^ mouse model was generated by Takaomi Saido and colleagues at the RIKEN Brain Science Institute in Japan [3]; the PS19 (B6;C3-Tg(Prnp-MAPT*P301S)PS19Vle/J, stock #008169) and C57BL/6J (stock #000664) mice were originally purchased from the Jackson Laboratory in Maine [24]. All mice used were on a congenic C57BL/6J background. To generate the APP^NL-G-F^/MAPT^P301S^ cross, homozygous APP^NL-G-F^ mice were bred with heterozygous PS19 mice resulting in either the double transgenic cross or heterozygous APP^NL-G-F^ mice. Due to littermates being heterozygous for the APP mutations, wild-type C57BL/6J mice were used as controls.

### Immunoblot

The homogenized lysate for western blot were collected from fresh frozen brain tissue with RIPA lysis buffer. Reducing and non-reducing protein samples were separated by gel electrophoresis and transferred to 0.2µm nitrocellulose membranes using the Bolt SDS-PAGE system (Life Technologies). Membranes were blocked in 5% nonfat dry milk (NFDM) in PBS supplemented with 0.025% Tween-20 (PBST) for 1 hour RT, followed by incubation overnight at 4°C in primary antibody diluted in 5% bovine serum albumin/PBST. Primary antibodies used were as follows: pTau217 (1:500) anti-tau antibody (rabbit, Thermo Scientific, Cat# 44744); 4G8, Anti-Amyloid β Antibody, clone W0-2, reactive to amyloid-β, aa 17-24 (Millipore Sigma, Cat# MABN10). Membranes were then washed 3 times with PBST and incubated in HRP-conjugated secondary antibodies (Jackson ImmunoResearch) diluted in 1% BSA/PBST at RT for 1 h. After incubation in secondary antibody, membranes were washed 3 times in PBST and developed using SuperSignal West Pico Chemilluminescent ECL substrate (ThermoFisher Scientific, cat# 34080).

### Immunohistochemistry

Wild type (WT), APP^NL-G-F,^ MAPT^P301S^, and APP^NL-G-F^/MAPT^P301S^ mouse brains were collected at 3, 6 and 9-month of age, respectively. Briefly, mice were anaesthetized with isoflurane and then the hearts perfused with 20 ml ice cold PBS for 5 mins followed by perfusion with 20ml ice cold 4% PFA for 10 mins. The mouse brains were dissected and placed in 4% PFA on ice for 2 hours. Then the brains were washed with PBS and transferred into 30% sucrose/PBS until the brains sank to the bottom of the tube (about 48h), and sectioned. The fixed brains were sliced into 30µm coronal sections by cryostat, and stored in 0.005% sodium azide/PBS solution at 4°C for up to 3 months. For long-term storage, the sections were transferred into cryoprotectant solution (30% glycerol and 30% ethylene glycol in PBS), and stored at -20 °C.

For immuno-labeling, the 30 µm free-floating sections with hippocampus or lateral entorhinal cortex (LEnt) were blocked with 5% BSA and 5% goat serum in PBST (PBS/0.25% Triton X-100) for 30 min and then incubated with monoclonal 6E10 antibody (BioLegend, cat# 803001, 1:1000 dilution) overnight at 4 °C. On the second day, sections were washed with PBST three times and then incubated with biotinylated goat anti-mouse IgG antibody (Vector Laboratories, cat# BA-9200) for 2 hours RT. The antibody binding was visualized using a Vectastain ABC Kit (Vector Laboratories, cat# PK-6100) and diaminobenzidine (DAB) substrate tablet (Sigma-Aldrich, cat# D4293-50SET) as described previously [25]. Images were captured by Keyence microscope bz-x800.

### Immuno-fluorescence staining of fixed brain tissues

For immuno-fluorescence labeling, selected sections of hippocampus from bregma -1.8 and LEnt from bregma -2.8, were washed in PBS for 10 mins and then permeabilized in 0.5ml PBS/0.25% Triton X-100 (PBST). Block tissues in blocking solution supplemented with 5% BSA and 5% normal donkey serum in PBST, 1.5-2 hours at RT. Dilute 1° antibodies in 5% BSA/PBST and incubate sections for overnight at 4°C. On the second day, wash the brain sections 3x in PBST, 15 min each. And then incubate the brain sections in 2° antibodies (1:700 for Dylight-/Alexa-conjugated antibodies made in donkey purchased from Thermo Fisher Scientific) in 5% BSA/PBST and incubate for 2 hours at RT. For DAPI nuclei stain, dilute DAPI (1:10,000) in PBST and incubate for 15 min. Wash 2x with PBST then 1x with PBS, 10 min each. Mount the brain sections onto microscope glass slides in Prolong gold anti-fade reagent. The primary antibodies used in this study are as follows: NeuN (chicken, Millipore, cat# ABN91), 1: 300; MC1 (mouse, provided by Peter Davies, Northwell), 1:100; TOMA2 (mouse, provided by Rakez Kayed, UTMB Galveston), 1:200 [26]; Images were captured by Carl Zeiss confocal LSM700.

### Measurements of pathological proteins Aβ and phosphorylated Tau by Enzyme-linked immunosorbent assay (ELISA)

The Measurements of pathological Aβ and phosphorylated tau by ELISA is as described previously [27-29]. In brief, frozen mouse brain tissue was homogenized in 5:1 volume of freshly prepared, ice cold 5M guanidine hydrochloride in Tris-buffered saline (20 mM Tris-HCl, 150 mM NaCl, pH 7.4), which contained 1:100 Halt protease inhibitor cocktail (Thermo Fisher Scientific) and 1:100 phosphatase inhibitor cocktail 2 & 3 (Sigma-Aldrich) as previously reported [27, 28]. The homogenate was then shaken (regular rocker) overnight at room temperature. The lysate was diluted with 1% Blocker A (Meso Scale Discovery [MSD], #R93BA-4) in wash buffer according to specific immunoassays: 1:4000 for Aβ_1-38_, Aβ_1-40_, and Aβ_1-42_ (MSD #K15200E-2), and 1:300 for p-tau_181_, p-tau_202_ (MSD custom kit), total tau and p-tau_231_ (MSD #K15121D-2). Samples were centrifuged at 17,000 g and 4°C for 15 minutes. The supernatant was subsequently applied to the immunoassays, and the original homogenate was aliquoted and stored at −80°C.

To capture MAPT phosphorylated at Thr residue 181, antibody AT270 was used. The detecting antibody was the biotinylated HT7 that recognizes residue 159-163 of tau (Thermo Fisher Scientific). To measure p-Tau^396^, a rabbit monoclonal antibody against p-Tau^396^ (Abcam, ab156623) was used as the capturing antibody, and HT7 was used as a detecting antibody. Sulfotag conjugated streptavidin secondary antibody was used for signal detection by the MSD platform. MSD SECTOR Imager 2400 was used to measure p-tau_396_ levels. Internal calibrators of p-Tau and tau were used (MSD). P-Tau levels were measured in arbitrary units, which may or may not be related among the different epitopes. Standards with known concentrations were used for Aβ, and all standards and samples were run in duplicate. Measurements were made using the multi-detection SPECTOR 2400 Imager (MSD).

### RT qPCR

The reverse transcription quantitative real-time PCR (RT-qPCR) was applied for the rapid detection of gene expression changes of pro-inflammatory cytokines and complement proteins in the brain tissue of WT, APP^NL-G-F^, MAPT^P301S^, and APP^NL-G-F^/MAPT^P301S^ double transgenic, respectively. The primers used in this study are listed below:

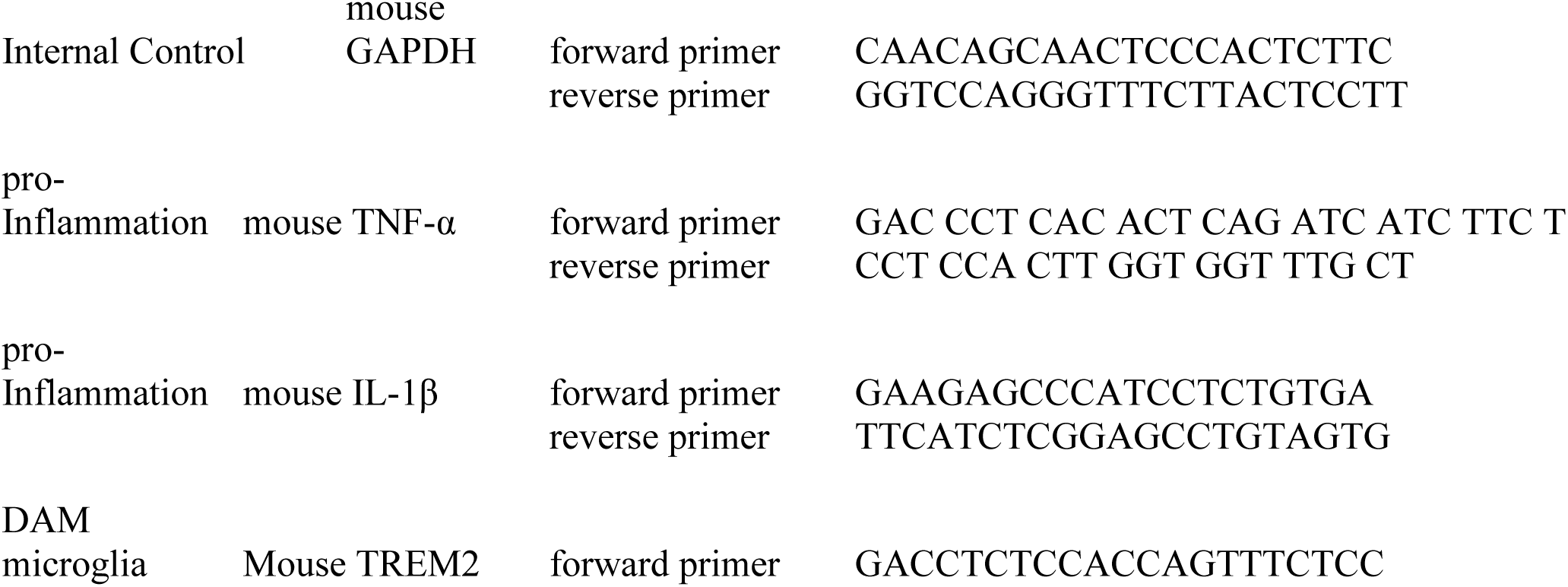

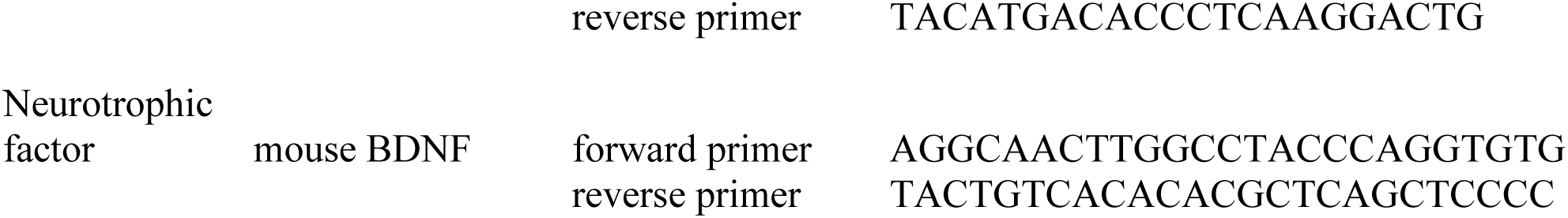

### Images Analysis

The intensity in immuno-fluorescence or DAB stained brain sections were measured by MATLAB program; the NeuN-positive cell number were quantified by Image J automatically cell counting. The quantification of cell numbers was done in a blinded manner, with the investigator analyzing the samples being different than the investigator coding the samples.

### Statistical analysis

Statistical analyses and figures artwork were performed using GraphPad Prism version 9.00 for Windows with two sided α of 0.05. All group data are expressed as mean ± SEM. Colum means were compared using one-way ANOVA with treatment as the independent variable. And group means were compared using two-way ANOVA with factors on genotype and age time course of the mice, respectively. When ANOVA showed a significant difference, pair wise comparisons between group means were examined by Tukey’s multiple comparison test. Significance was defined at *p*< 0.05.

## Results

### Generation of the APP^NL-G-F^/MAPT^P301S^ mouse line

The APP^NL-G-F^ mouse line was developed to avoid artifacts associated with over-expressing APP [3]. These mice develop robust neuritic plaque pathology. We initiated the project by crossing homozygous APP^NL-G-F^ mice with heterozygous P301S MAPT mice. The mice generated normal mendelian patterns of inheritance, producing expected genotypes in the offspring. The resulting mouse lines were aged, harvested and examined patterns of APP expression, as well as the accumulation of Aβ and neuritic plaques at 3, 6 and 9 months of age. Immunoblots of APP using the 48G antibody showed that all mice exhibited similar levels of APP expression at each age (**Fig. 1A, B**).

**Figure 1.**
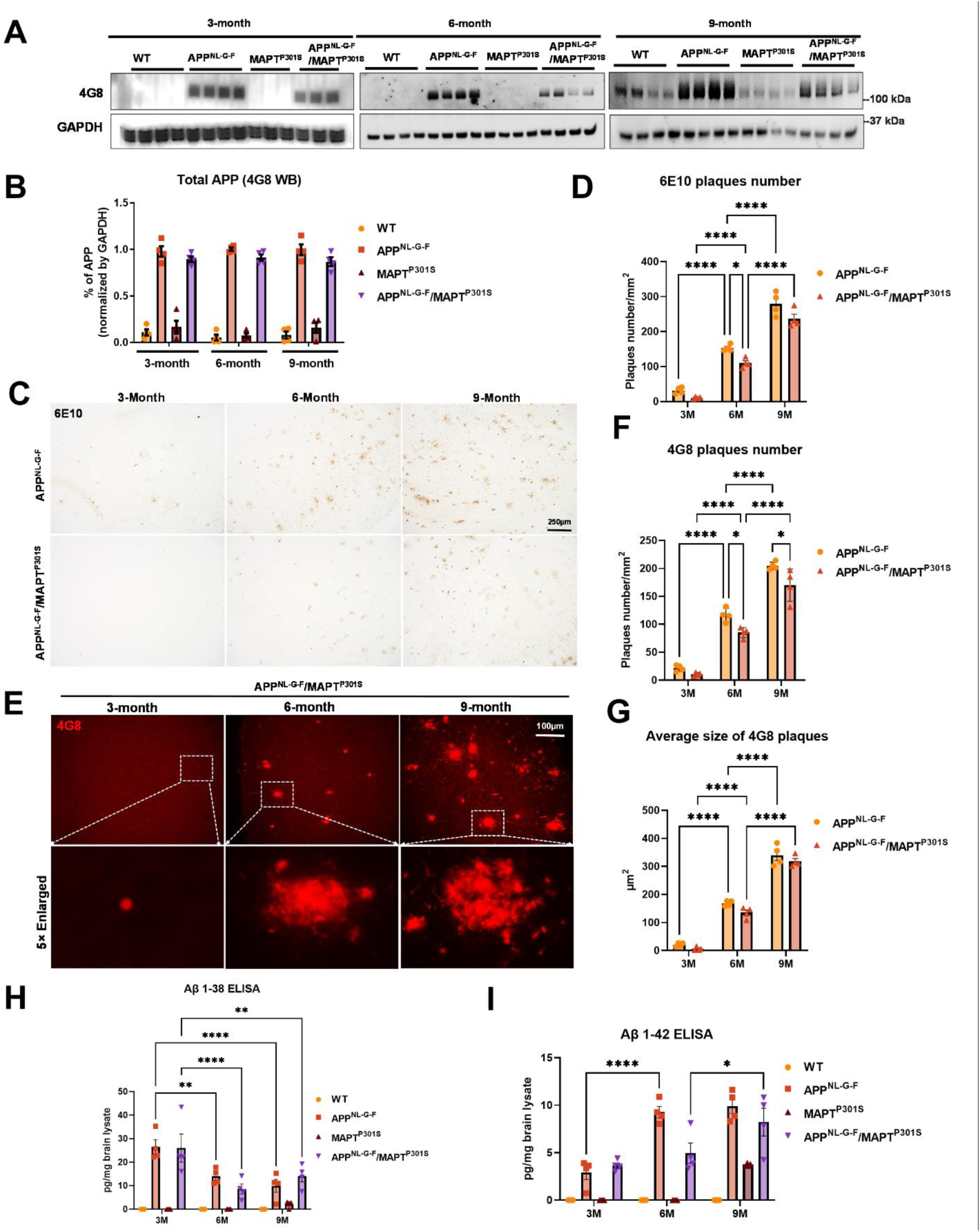
Beta-Amyloid deposition accumulates in a time-dependent manner in the APP^NL-G-F^/MAPT^P301S^ mouse. **A.** Representative images of western blot with 4G8 antibody showed the expression of human amyloid precursor protein (APP) in the APP^NL-G-F^ and APP^NL-G-F^/MAPT^P301S^ mouse brain but not wild type (WT) or MAPT^P301S^ brain. Total brain lysates were harvested at 3, 6 and 9 month old in each of the four mouse genotypes (WT, APP^NL-G-F^, MAPT^P301S^, and APP^NL-G-F^/MAPT^P301S^ double transgenic, respectively). GAPDH was detected as the internal control. **B.** Quantification of human APP expression in the total brain lysate as shown in (A). N=4, Data shown as mean ± SEM. **C.** The 6E10 antibody (reactive to aa 1-16 Aβ and to APP) was used to examine the diffused amyloid plaques in the aging process of APP^NL-G-F^ and APP^NL-G-F^/MAPT^P301S^ mouse brain. Representative DAB staining images showed the progressive increase of 6E10 positive β-amyloid plaques in the entorhinal cortex from 3-month to 6 and 9-month of mouse brain. Scale bar 250µm. **D.** Quantification for the number of 6E10 positive β-amyloid plaques averaged over 1 mm^2^ squares across each brain slice. N=4 mice in each group, Data shown as mean ± SEM. Two-way ANOVA with Tukey’s multiple comparisons test, *p<0.05, ****p<0.001. **E.** The 4G8 antibody (reactive to Aβ, aa 17-24) was used to examine the compact amyloid plaques in the aging process of APP^NL-G-F^ and APP^NL-G-F^/MAPT^P301S^ mouse brain. Representative red fluorescence labeling stacked images showed the progressive increase of 4G8 positive β-amyloid plaques in the entorhinal cortex from 3-month to 6 and 9-month of mouse brain. Scale bar 100µm. **F-G**. Quantification of the number and average size of 4G8 positive Aβ+ plaques. N=4 mice per group, 3 sections were used for each mouse. Data shown as mean ± SEM. Two-way ANOVA with Tukey’s multiple comparisons test, *p<0.05, ****p<0.001. **H-I**. The amount of Aβ_38_ and Aβ_42_ in the total brain lysate detected by V-PLEX Aβ Peptide Panel 1 (4G8) Kit. Brain lysate from 4 genotypes of mice were detected at 3, 6 and 9 months, respectively. N=4 mice in each condition, data shown as mean ± SEM. Two-way ANOVA with Tukey’s multiple comparisons test, *p<0.05, **p<0.01, ****p<0.001.

Next we explored whether the accumulation of Aβ was affected by the presence of the P301S MAPT transgene. Mice at 3, 6 and 9 months were harvested and subjected to immunohistochemistry using both colorimetric and fluorescent approaches. DAB/peroxidase colorimetry can capture the broad distribution of pathology and also avoids any complications from lipofuscin accumulation. Immunofluorescence was also used because it can provide quantification based on intensity and provides finer imaging detail. Analysis of sections with antibody 6E10, which preferentially detects diffuse plaques, was done using the colorimetric DAB/peroxidase method. These results showed a progressive increase in amyloid plaque accumulation in the APP^NL-G-F^ mouse line as well as the APP^NL-G-F^/MAPT^P301S^ mouse line (**Fig. 1C, D**). The presence of the MAPT^P301S^ exerted only a small effect on the accumulation of 6E10 positive plaques, with a 20% decrease in plaque load evident only at 6 months (p<0.05, Fig. 1C, D). Studies with 4G8, which preferentially detects consolidated plaques, showed the same pattern (**Fig. 1E-G**). A progressive increase in plaque load was observed in both the APP^NL-G-F^ mouse line as well as the APP^NL-G-F^ x MAPT^P301S^ mouse line; we did note that the APP^NL-G-F^ x MAPT^P301S^ mice exhibited slightly fewer (15-20%) plaques than the APP^NL-G-F^ mice at 6 and 9 months (**Fig. 1F**), however the average plaque size did not differ between the two groups (**Fig. 1G**).

To explore the levels of extracellular Aβ and neuritic plaques in the transgenic mouse, we examined the amount of Aβ_38_ and Aβ_42_ in the total brain lysate by V-PLEX Aβ Peptide ELISA Kit. The V-PLEX platform offers analytically validated singleplex and multiplex assay kits, which can provide accurate and reproducible results with consistency from lot to lot [30]. The ELISA result showed that Aβ_38_ levels were similar in APP^NL-G-F^ and APP^NL-G-F^ x MAPT^P301S^ mice at 3 month old but slowly decreased as the mice aged. In contrast, the Aβ_42_ level was elevated and progressively increased with age in both APP^NL-G-F^ and APP^NL-G-F^ x MAPT^P301S^ mouse brains. Notably, in the APP^NL-G-F^ mouse the Aβ_42_ level peaked at 6-months and remained constant at 9-months while levels of Aβ_42_ steadily increased with age (at 3, 6 and 9-months) in APP^NL-G-F^ x MAPT^P301S^ mouse brain.

These results indicate that the APP^NL-G-F^ x MAPT^P301S^ double transgenic mouse model recapitulates the progressive accumulation Aβ plaques and levels characteristic of brains of AD subjects.

### APP^NL-G-F^ potentiates progression of MAPT pathology in the APP^NL-G-F^/MAPT^P301S^ double transgenic mice

Another goal of the APP^NL-G-F^/MAPT^P301S^ double transgenic mice is to recapitulate the development of MAPT pathology associated with cognitive decline in AD patients. To investigate the MAPT aggregation in the MAPT^P301S^ and APP^NL-G-F^/MAPT^P301S^ mice as well as the effect of APP^NL-G-F^ on MAPT pathology, we examined MAPT phosphorylation, oligomerization and misfolding in the aging process of all four genotypes, including MAPT^P301S^, APP^NL-G-F^ and APP^NL-G-F^/MAPT^P301S^ in comparison to WT C57BL/6 control. The total brain lysates were prepared from the fresh frozen brain tissue harvested at 3, 6 and 9-months and homogenized in RIPA buffer.

Recent studies highlight phosphorylated MAPT at threonine MAPT 217 (pTau217) as a new promising plasma biomarker for pathological changes implicated in AD [31]. Immunohistochemistry with postmortem AD brain tissue also demonstrated that pTau217 is found in neurofibrillary tangles (NFTs) and neuropil threads that are also positive for pTau181, 202, 202/205, 231, and 369/404 [32]. Levels of pTau217 also correlate with total Aβ and NFT brain load in AD brain [32, 33]. Thus, we use levels of pTau217 as biomarker for evaluating the pathological progression of the APP^NL-G-F^/MAPT^P301S^ mouse model. Western blot of brain lysates from MAPT^P301S^ and APP^NL-G-F^/MAPT^P301S^ mice showed progressive accumulation of pTau217 over 3, 6 and 9-month range (**Fig. 2A, B**). The APP^NL-G-F^/MAPT^P301S^ exhibited significantly more pTau217 MAPT phosphorylation at 6 and 9-months compared to MAPT^P301S^ alone, which suggest that APP^NL-G-F^ potentiated MAPT phosphorylation (**Fig. 2A, sB**). To further confirm the accumulation of hyper-phosphorylated MAPT in the brain lysate and exclude the variation on the total MAPT expression level, we quantified the MAPT phosphorylation at threonine181 (pTau181) with the AT270 antibody and total tau level with BT2 antibody (epitope between aa 194-198, but not PHF MAPT), respectively, by ELISA assay. The result showed that APP^NL-G-F^/MAPT^P301S^ and MAPT^P301S^ alone mice presented similar levels of total MAPT but APP^NL-G-F^/MAPT^P301S^ contained significant elevated level of pTau181 (**Fig. 2C-D**).

**Figure 2.**
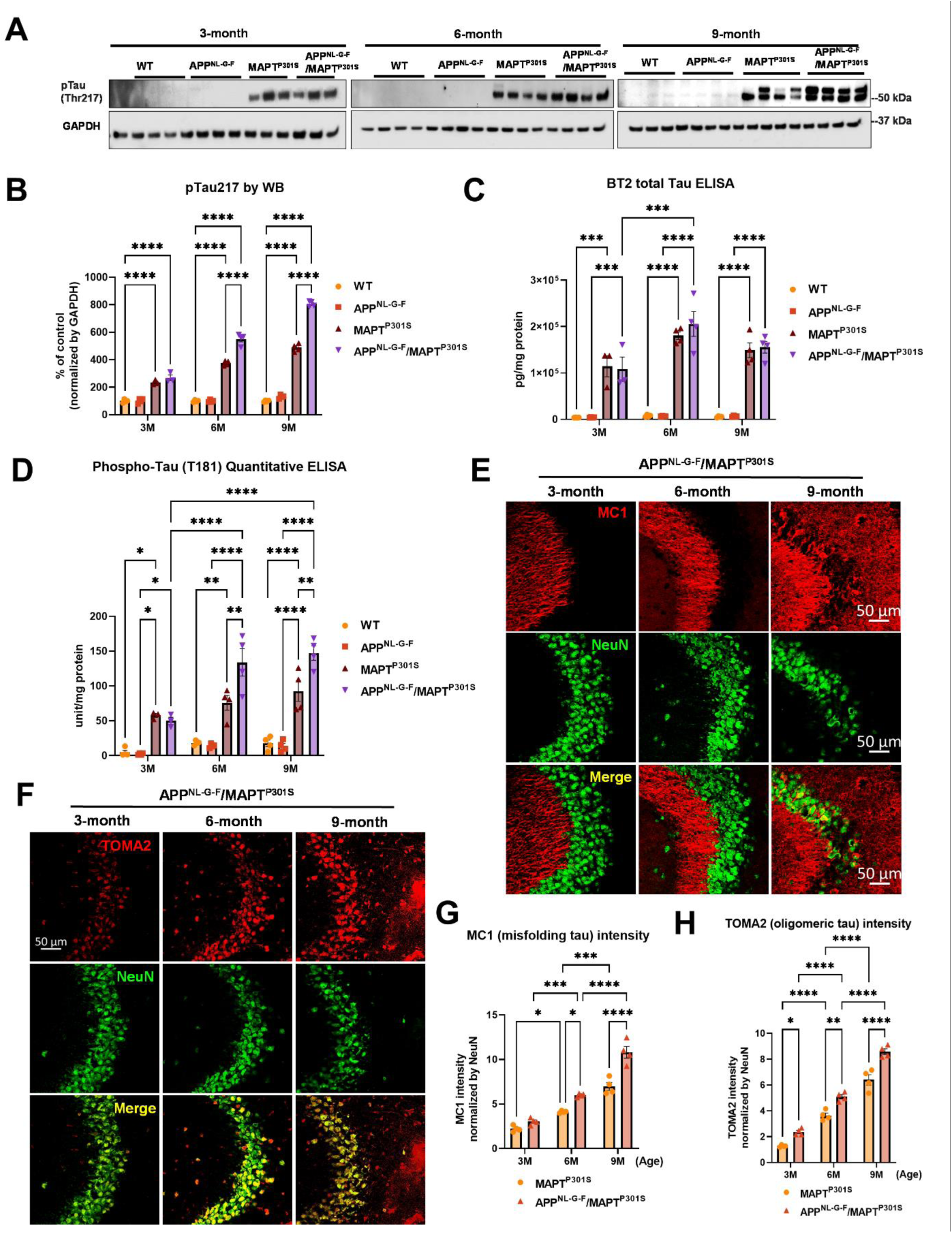
APP^NL-G-F^ potentiates progression of MAPT pathology in the APP^NLGF^/MAPT^P301S^ double transgenic mice. **A.** Representative images of western blot with phosphorylated tau antibody on phosphor-site threonine217 (pTau217) showed the accumulation of hyper phosphorylated tau in the MAPT^P301S^ and APP^NL-G-F^/MAPT^P301S^ mouse brain but not wild type (WT) or APP^NL-G-F^ brain. Total brain lysate were harvested at 3, 6 and 9 months for each of the four genotypes (WT, APP^NL-G-F^, MAPT^P301S^, and APP^NL-G-F^/MAPT^P301S^) of mice, respectively. GAPDH was detected as the internal control. **B.** Quantification of pTau217 in the total brain lysate as shown in (**A**). N=4 mice in each group, Data shown as mean ± SEM. Statistics was by two-way ANOVA with *post hoc* Tukey’s multiple comparisons test, ****p<0.001. **C-D**. Detection of total MAPT levels in the brain lysates with the BT2 antibody (epitope between aa 194-198, but not PHF tau) and threonine181 phosphorylated tau (pTau181) by AT270 antibody, respectively, with ELISA assay. N=4 mice in each group, Data shown as mean ± SEM. Statistical analysis was by two-way ANOVA with *post hoc* Tukey’s multiple comparisons test, ***p<0.005, ****p<0.001. **E.** Representative fluorescence labeling images showed the accumulation of misfolding tau (by MC1 antibody, red) in the hippocampal CA3 region of APP^NL-G-F^/MAPT^P301S^ mice over 3, 6 and 9 months. NeuN antibody (green) was used to label the neuronal cells. Scale bar 50µm. **F.** Representative fluorescence images show the accumulation of oligomeric tau (TOMA2 antibody, red) in the hippocampal CA3 brain region of APP^NL-G-F^/MAPT^P301S^ mice over 3, 6 and 9 months. NeuN antibody (green) was used to label neurons. Scale bar 50µm. **G-H**. Quantification of misfolded tau (MC1) and oligomeric tau (TOMA2) as shown in (**E**) and (**F**) respectively. Total fluorescence intensity was collected and then normalized by NeuN for statistics. N=4 mice in each group, data is shown as mean ± SEM. Two-way ANOVA was used for statistics followed by post hoc analysis with Tukey’s multiple comparisons test, *p<0.05, **p<0.01, ***p<0.005, ****p<0.001.

In addition to the MAPT phosphorylation, we also examined the MAPT aggregation by the conformational tau marker MC1 (epitope within aa 312-322) with immuno-fluorescence labeling. Our result showed that MAPT tau misfolding started at the dendritic compartment of the neurons in CA3 when the mice were 3-month old. The misfolded MAPT distributed and accumulated more in neuronal soma than dendrites and synapses when the mice were 9 month old (**Fig 2E**). Compared to MAPT^P301S^ alone, the APP^NL-G-F^/MAPT^P301S^ double transgenic mouse presented accelerated and potentiated accumulation of misfolded MAPT over the time span of 3, 6 and 9 month of age (**Fig. 2G**).

Studies suggest that MAPT oligomers are the more toxic species that induce neurodegeneration [25, 26, 34-36]. To assess the assembly of toxic MAPT oligomers in the APP^NL-G-F^/MAPT^P301S^ double transgenic mouse, we labeled MAPT with the antibody TOMA2, which is the oligomeric MAPT specific by immuno-fluorescence labeling in the hippocampus. The result showed that oligomeric MAPT accumulated in the somatic compartment of the neurons and was elevated in older animals (9 m) (**Fig. 2F**). Quantification of TOMA2 intensity revealed that MAPT oligomers were more abundant in APP^NL-G-F^/MAPT^P301S^ double transgenic mouse compared to MAPT^P301S^ (**Fig. 2H**).

These results demonstrate that APP^NL-G-F^ potentiates the progression of MAPT pathology including phosphorylation, mis-conformation and oligomerization in the APP^NL-G-F^/MAPT^P301S^ double transgenic mice.

### APP^NL-G-F^ is the predominant driver of microglial activation and astrogliosis

In addition to Aβ plaques and MAPT pathology, neuroinflammation including microglial activation and astrogliosis are thought to be key drivers mediating neurodegeneration in AD [37, 38]. To characterize the glial activation feature in the APP^NL-G-F^/MAPT^P301S^ double transgenic mouse, we analyzed the microglial and astrocytic morphology changes as well as the transcriptomic level of inflammatory factors in the aging process of APP^NL-G-F^/MAPT^P301S^ mouse brain. Our data showed that microglia exhibit a ramified appearance under basal conditions labeled by Iba-1 marker. The microglia then become hyper-ramified with elongation of their processes and increase branch complexity in MAPT^P301S^ mouse brain, suggestive of an intermediate step to microglial activation or a mild response. In the APP^NL-G-F^ and APP^NL-G-F^/MAPT^P301S^ mouse brain, microglia bcome highly activated displaying morphological changes with thickening processes and amoeboid shape (**Fig. 3A-B**). At the same time, astrocytes (labeled by astrocytic marker GFAP) are robustly activated in APP^NL-G-F^ and APP^NL-G-F^/MAPT^P301S^ mouse brains exhibiting enlarged morphological sizes and co-localization with the Aβ plaques (**Fig. 3A**). The morphological changes of microglia and astrocyte are particularly enhanced in the conditions with APP^NL-G-F^, implying that APP^NL-G-F^ is the predominant driver of microglial activation and astrogliosis (**Fig. 3C-D**).

**Figure 3.**
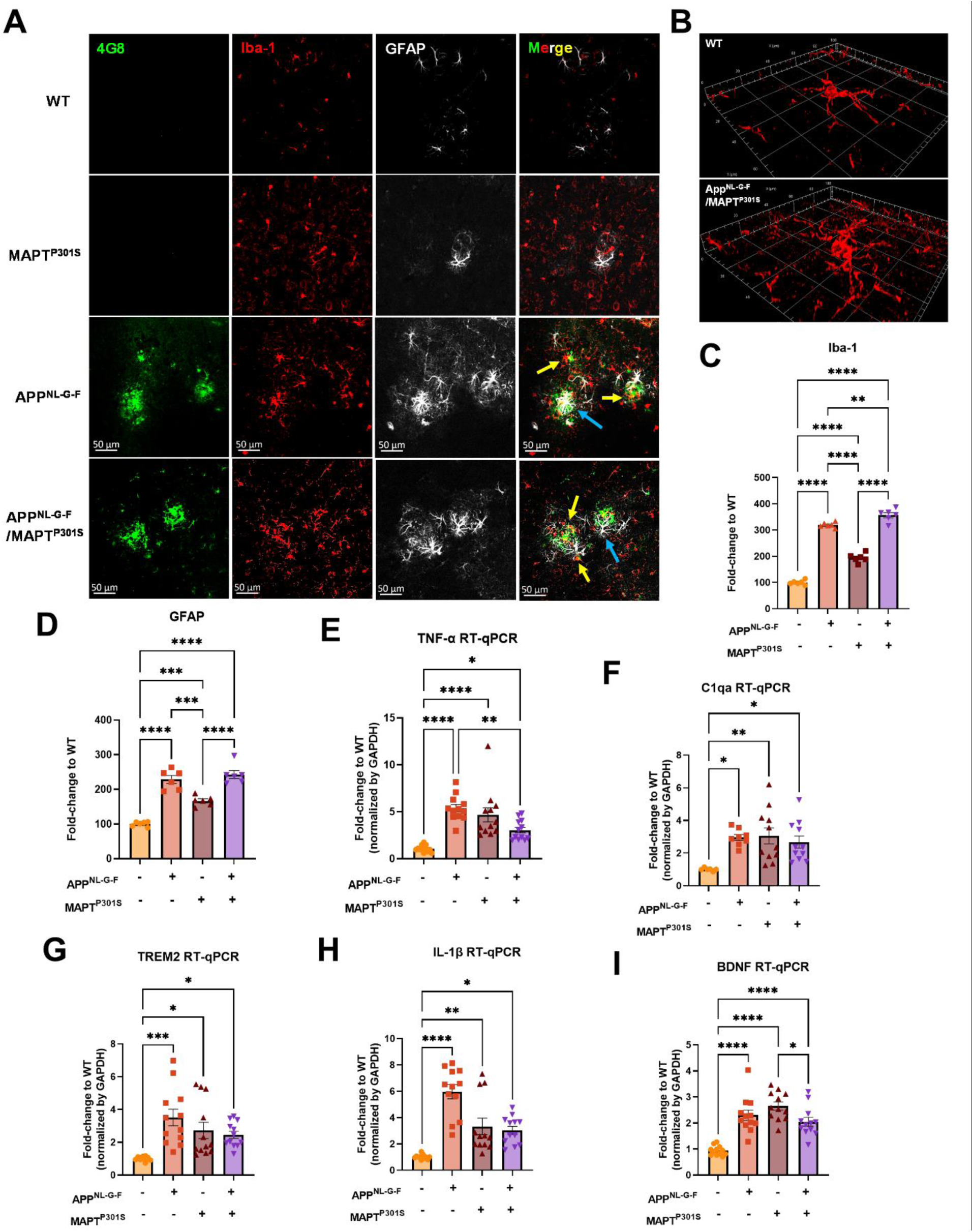
APP^NL-G-F^ is the predominant driver of microglial activation and astrogliosis. **A.** Representative fluorescence labeling images showed the activation and morphological changes of microglia (by Iba-1 antibody, red) in the frontal cortex in pathological APP^NL-G-F^ and/or MAPT^P301S^ mouse brain at 9-month old. Astrocytes (GFAP antibody, white) were robustly activated around Aβ plaques. Scale bar 50µm. **B**. Enlarged image showed the amoeba-like morphological changes of microglia in the APP^NL-G-F^/MAPT^P301S^ mouse brain. **C-D**. Quantification of microglial activation by Iba-1 intensity and astrogliosis by GFAP intensity as shown in (**A**). Data was normalized to the fold increase of WT control. N=6 mice in each group, data is shown as mean ± SEM. Two-way ANOVA was used for statistics followed by post hoc analysis with Tukey’s multiple comparisons test, **p<0.01, ***p<0.005, ****p<0.001. **E-I**. Quantification on the transcriptomic levels of inflammatory factors in the brain of WT, APP^NL-G-F^, MAPT^P301S^, and APP^NL-G-F^/MAPT^P301S^ mouse lines, respectively, at 9-months. The pro-inflammatory factors TNF-α, IL-1β, C1qa as well as BDNF were quantified by RT qPCR. Results are shown as fold-change vs. WT control. N=10-12 mice per group; data is shown as mean ± SEM. One-way ANOVA was used for statistics followed by post hoc analysis with Tukey’s multiple comparisons test, *p<0.05, **p<0.01, ***p<0.005, ****p<0.001.

To further confirm the microglia and astrocyte activation, we measured transcripts coding for pro-inflammatory cytokines, anti-inflammatory cytokines and complement proteins in the brain tissue of WT, APP^NL-G-F^, MAPT^P301S^, and APP^NL-G-F^/MAPT^P301S^ mice, respectively. Our result show that the pro-inflammatory cytokine TNF-α and IL-1β are increased in all APP^NL-G-F^, MAPT^P301S^ and APP^NL-G-F^/MAPT^P301S^ mice compared to WT mice. Interestingly, the APP^NL-G-F^ mouse brain showed the highest levels of the two inflammatory factors (**Fig. 3E, H**). The complement protein, C1q, has been shown to bind to fibrillar Aβ and NFTs in AD resulting in the activation of the classical complement pathway [39, 40]. In the APP^NL-G-F^/MAPT^P301S^ mouse model, we examined the C1qa RNA level by RT-qPCR and found that it was elevated equally among the APP^NL-G-F^, MAPT^P301S^, and APP^NL-G-F^/MAPT^P301S^ mouse lines compared to WT control (**Fig. 3F**). Next we analyzed TREM2, a key player in microglial biology and AD[41]. Analysis of TREM2 transcripts demonstrated a similar (∼3-fold) increase in the APP^NL-G-F^, MAPT^P301S^, and APP^NL-G-F^/MAPT^P301S^ mouse lines (**Fig. 3G**). Finally, we also examined brain-derived neurotrophic factor (BDNFs), which maintains synaptic plasticity and has attracted increasing attention for its potential as a biomarker or therapeutic molecule for AD [42]. Quantification of BDNF transcript by RT-PCR in the APP^NL-G-F^/MAPT^P301S^ mouse model showed elevation BDNF each of the mouse models(**Fig. 3I**).

The combined data indicate that the APP^NL-G-F^/MAPT^P301S^ mouse model recapitulates many of the pivotal neuroinflammatory features of AD, and also suggest that the accumulation of Aβ is a stronger driver of microglial activation and astrogliosis that MAPT pathology.

### APP^NL-G-F^ potentiates neurodegeneration in the APP^NL-G-F^/MAPT^P301S^ mice

In the current APP^NL-G-F^/MAPT^P301S^ mouse model, we have shown multiple hallmark features of AD, including Aβ plaques, MAPT pathology and neuroinflammation. To further understand their roles in mediating neurodegeneration, we quantified how expression of neuronal and synaptic markers change with aging in each mouse line (APP^NL-G-F^, MAPT^P301S^, and APP^NL-G-F^/MAPT^P301S^ vs. WT control). Our results showed that the APP^NL-G-F^/MAPT^P301S^ mouse line displayed progressive neuronal loss in consistency with Aβ plaques and tau pathology in the aging process from 3 to 6 and 9 month old (**Fig. 4A**). At 6-months, the APP^NL-G-F^/MAPT^P301S^ mouse showed ∼35% neuronal loss compared to WT control, which progressed to more than 60% loss of MAP-2 at 9-months (**Fig. 4B**). We aimed to elucidate the contribution of APP^NL-G-F^ and MAPT^P301S^ in neuronal loss by comparing the APP^NL-G-F^/MAPT^P301S^ mouse to APP^NL-G-F^ KI alone and P301S Tau PS19 conditions. Our result showed that APP^NL-G-F^ KI itself exhibited limited amount of neuronal loss but can potentiate the neuronal loss in APP^NL-G-F^/MAPT^P301S^ mouse brain in comparison to P301S MAPT PS19 over-expression alone (**Fig. 4C-D**). In addition to the immuno-fluorescence labeling, we also applied the biochemistry assay with post-synaptic marker PSD-95 probed by western blot to validate the neuronal deficit in APP^NL-G-F^/MAPT^P301S^ mouse brain (**Fig. 4E**). The quantification of PSD-95 showed that MAPT^P301S^ is correlated to the neuronal deficit while APP^NL-G-F^ can potentiate the neuronal loss mediated by MAPT^P301S^ (**Fig. 4F**).

**Figure 4.**
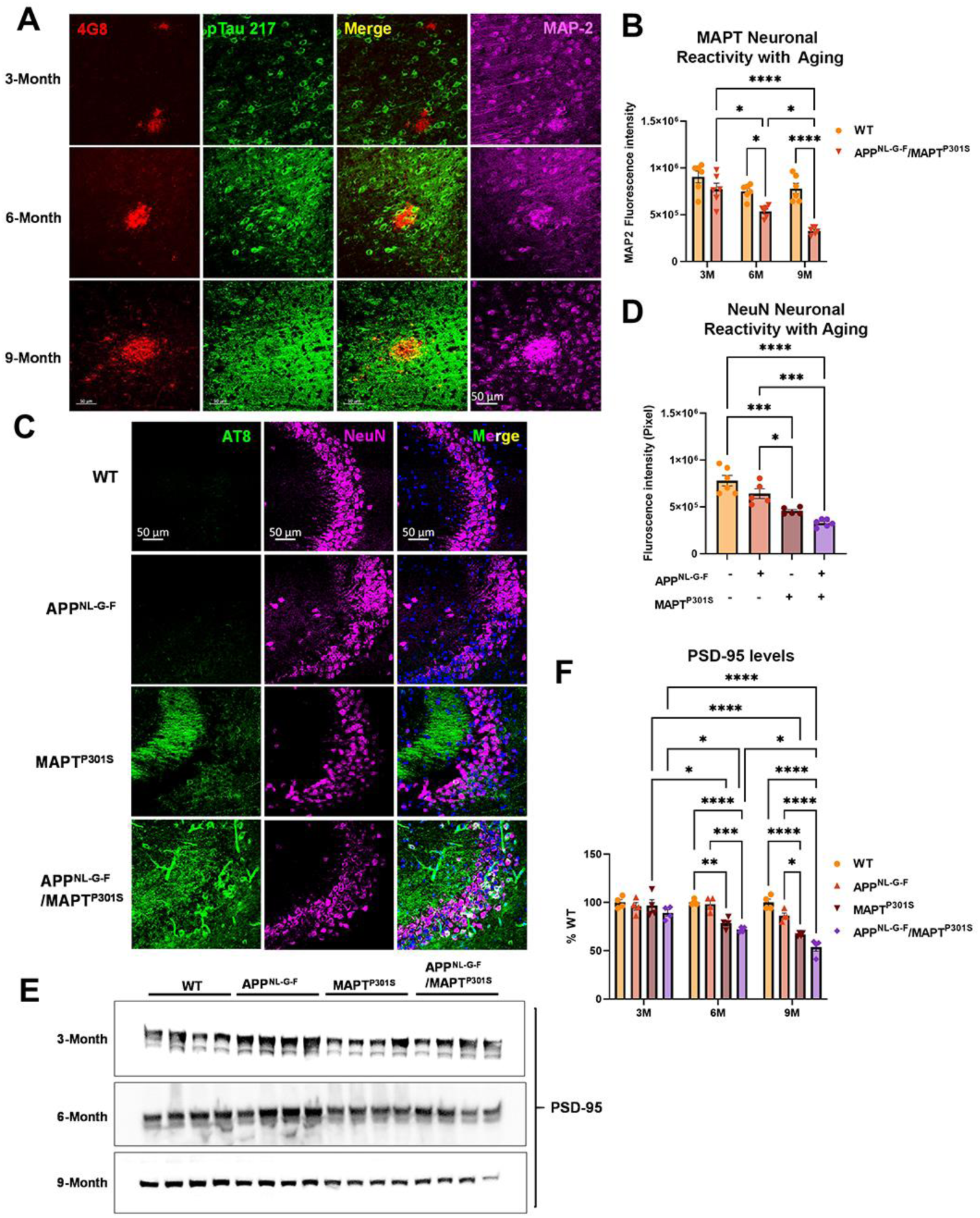
APP^NL-G-F^ potentiates neurodegeneration in the APP^NLGF^/MAPT^P301S^ double transgenic mice. **A.** Representative images showed the enhanced neurodegeneration (MAP-2, magenta) associated with progressive Aβ deposition (4G8, red) and phosphorylated tau (pTau217, green) accumulation in the APP^NL-G-F^/MAPT^P301S^ mouse brain. Scale bar 50µm. **B**. Quantification of neurodegeneration by MAP-2 intensity as shown in (**A**). N=6 mice in each group; data is shown as mean ± SEM. Two-way ANOVA was used for statistics followed by post hoc analysis with Tukey’s multiple comparisons test, *p<0.05, ****p<0.001. **C.** Representative images showed enhanced neurodegeneration in APP^NL-G-F^/MAPT^P301S^ mouse brain compared to APP^NL-G-F^ or MAPT^P301S^ mouse lines at 9-months. Scale bar 50µm. **D.** Quantification of neurodegeneration by NeuN positive neuronal intensity as shown in panel C (magenta panels). N=6 mice in each group, data is shown as mean ± SEM. One-way ANOVA was used for statistics followed by post hoc analysis with Tukey’s multiple comparisons test, *p<0.05, ***p<0.005. **E-F**. Western blot of post-synaptic marker PSD-95 showed the potentiated neurodegeneration in the APP^NL-G-F^/MAPT^P301S^ mouse brain compared to APP^NL-G-F^ or MAPT^P301S^ mouse lines over a 3, 6 and 9-month old. Quantification of band intensities showed that progressive and enhanced decrease of PSD-95 in the APP^NL-G-F^/MAPT^P301S^ mouse brain. N=4 mice in each group, data was normalized to percentage of WT control and is shown as mean ± SEM. Two-way ANOVA was used for statistics followed by post hoc analysis with Tukey’s multiple comparisons test, *p<0.05, **p<0.01, ***p<0.005, ****p<0.001.

### N^⁶^-Methyladenosine and its regulator enzyme proteins are dysregulated in the APP^NLGF^/MAPT^P301S^ double transgenic mice in correspondence to the progression of tau pathology

N6-methyladenosine (m^6^A) is the most prevalent, abundant and conserved internal cotranscriptional modification in eukaryotic RNA [43]. In the recent studies, our group discovered that m6A accumulation is a general feature of AD pathology [23]. Levels of m^6^A modifications are controlled by addition of m^6^A modifications the m6A methyltransferases (also known as writers), such as METTL3/14/16, RBM15/15B and WTAP; m^6^A levels are also controlled by removal via demethylases (also known as erasers), including FTO and ALKBH5. The m^6^A-binding proteins YTHDF1/2/3, YTHDC1/2 IGF2BP1/2/3 and HNRNPA2B1 recognize the modifications; these are also known as “readers”[43]. Previously we showed that interaction of MAPT with HNRNPA2B1 and m^6^A RNA mediates the progression of tauopathy [23]. Other studies of AD indicate that m^6^A dysregulation often occurs in the context of altered expression of m6A writers and readers [44-48]. Hence we examined levels of m6A writers and readers in the APP^NL-G-F^/MAPT^P301S^ mouse model. Levels of m^6^A progressively increased in the APP^NL-G-F^/MAPT^P301S^ mouse at 3, 6 and 9-months compared to WT control (**Fig. 5A-B**). The m^6^A methyltransferase Mettl3 also increased in the APP^NL-G-F^, MAPT^P301S^ and APP^NLGF^/MAPT^P301S^ compared to WT control at 6-months. However, at 9 months only the MAPT^P301S^ or APP^NLGF^/MAPT^P301S^ mouse lines showed progressive increases in Mettl3 levels (**Fig. 5C-D**). The m^6^A eraser ALKBH5 showed a corresponding decrease in expression in MAPT^P301S^ and APP^NLGF^/MAPT^P301S^ mouse compared to WT control at 6 and 9 months (**Fig. 5E-F**). These results demonstrate dysregulation of m^6^A and enzymes in the m^6^A pathway in a manner similar to AD, and suggest that the phenomenon is predominantly driven by MAPT^P301S^ tau pathology.

**Figure 5.**
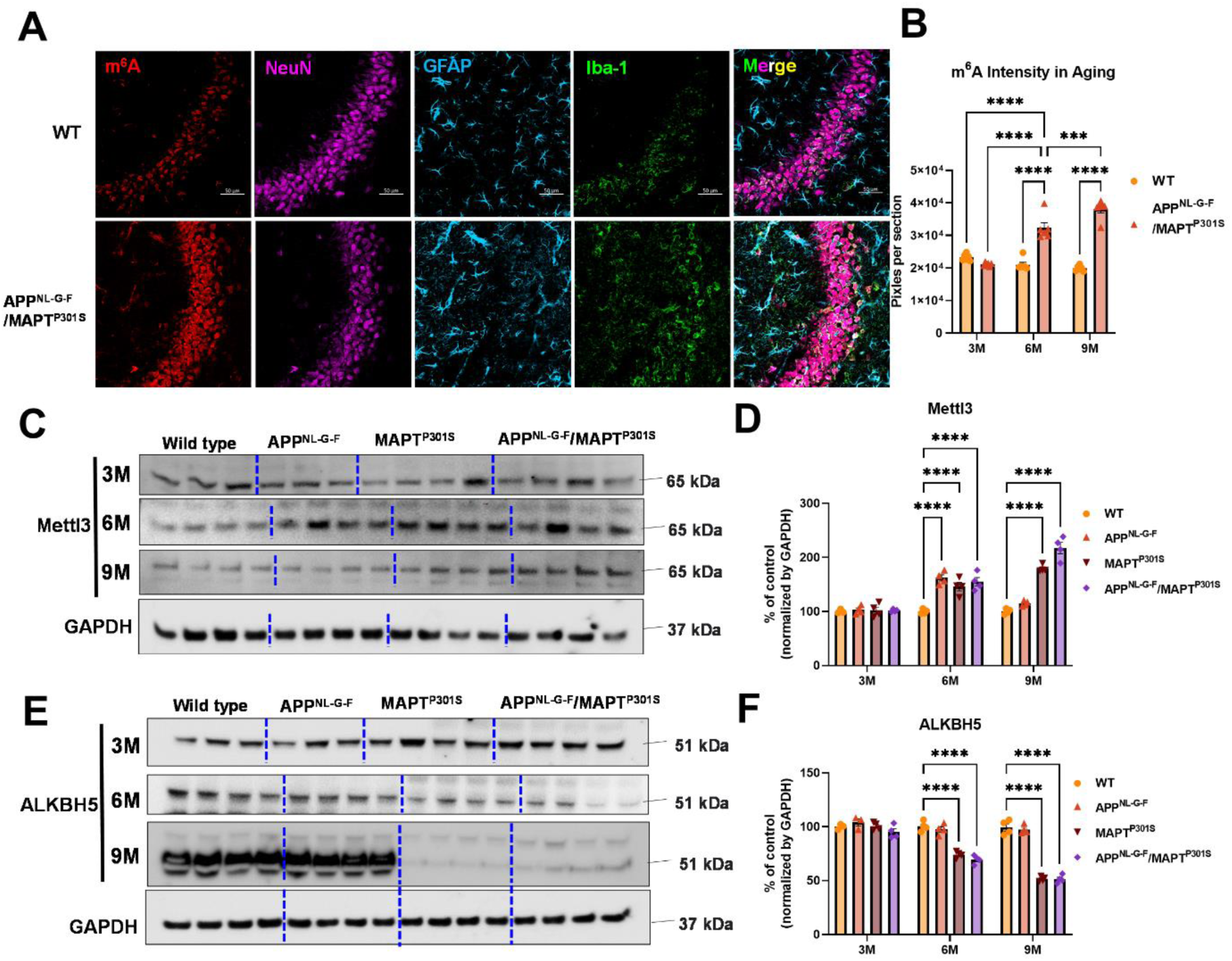
m^6^A and its regulator enzyme proteins are dysregulated in the APP^NLGF^/MAPT^P301S^ double transgenic mice in correspondence to the progression of tau pathology. **A.** Representative images showed the increased m^6^A intensity in APP^NLGF^/MAPT^P301S^ mouse brain at 6-months compared to WT control. Scale bar 50µm. **B.** Quantification of m^6^A intensity in comparison between APP^NLGF^/MAPT^P301S^ and WT control during the aging process. N=6 mice in each group, 3 brain sections were selected from each mouse brain with same position of hippocampus CA3. Data are shown as mean ± SEM. Two-way ANOVA was used for statistics followed by post hoc analysis with Tukey’s multiple comparisons test, ***p<0.005, ****p<0.001. **C-D**. Western blot analysis of the m^6^A methyltransferase Mettl3 showed progressively increased intensity in APP^NL-G-F^/MAPT^P301S^ mouse brain compared to APP^NL-G-F^ or MAPT^P301S^ alone at 3, 6 and 9 months. GAPDH was detected and used as internal control for statistical analysis. N=4 mice in each group, data was normalized to percentage of WT control and is shown as mean ± SEM. Two-way ANOVA was used followed by post hoc analysis with Tukey’s multiple comparisons test, ****p<0.001. **E-F**. Western blot analysis of the m^6^A RNA demethylase ALKBH5 showed decreased intensity in APP^NL-G-F^/MAPT^P301S^ mouse brain compared to APP^NL-G-F^ or MAPT^P301S^ alone during the aging process at 3 6 and 9 months. Quantification of band intensity showed the dereased ALKBH5 correlated with MAPT pathology in APP^NL-G-F^/MAPT^P301S^ and MAPT^P301S^ mouse brain. GAPDH was used as internal control for statistical analysis. N=4 mice in each group, data was normalized to percentage of WT control and is shown as mean ± SEM. Two-way ANOVA was used followed by post hoc analysis with Tukey’s multiple comparisons test, ****p<0.001.

## Discussion

The results presented above show that Aβ enhances tau pathologies (phosphorylation, misfolding, and fibrillization) in the context of P301S MAPT over-expression. Neurodegeneration parallels tau pathology, being enhanced in the double transgenic mouse. However, the converse is not true. Aβ pathologies (neuritic plaque, Aβ_40_ and Aβ_42_ load) are not greater in the APP^NL-G-F^/MAPT^P301S^ mouse than in the APP^NL-G-F^ mouse alone. These results are consistent with those recently reported for an APP^NL-G-F^/MAPT^P209L^ mouse [49]. The report on the APP^NL-G-F^/MAPT^P209L^ mouse showed enhancement of MAPT pathology by Aβ but did not provide any information on other pathologies, such as inflammation or m^6^A. We now report that the amount of inflammation parallels Aβ pathology, while the amount of m^6^A parallels MAPT pathology.

These results are generally consistent with prior observations using crosses of APP over-expression mouse lines and MAPT over-expression mouse models [12-17]. These over-expression models consistently observe that the presence of Aβ pathology enhances the accumulation of MAPT pathology, neurodegeneration and cognitive loss [12-17]. These same studies indicate that the presence of MAPT pathology either doesn’t change the accumulation of Aβ pathology or decreases it [12-17], with only one initial study suggesting that MAPT pathology increases Aβ pathology [18]. A closer look at the MAPT and Aβ pathologies suggest further parallels between the APP^NL-G-F^/MAPT^P301S^ cross and AD in humans. Bulbous processes filled with MAPT can be seen surrounding Aβ plaques in both the APP^NL-G-F^/MAPT^P301S^ cross and in AD. These bulbous processes are thought to reflect degenerating neuronal dendritic processes (Fig. 4A) [50]. The enhancement of MAPT pathology in the APP^NL-G-F^/MAPT^P301S^ mouse is also evident in the distribution of MAPT pathology in the hippocampus. The APP^NL-G-F^ model exhibits modest phospho-MAPT pathology at 6 and 9 months that is evident in the dendritic fields in the CA3 region, while the APP^NL-G-F^/MAPT^P301S^ cross exhibits MAPT pathology in the dendritic fields as well as in the neuronal soma of CA3 (Fig. 4D). The cell body MAPT pathology is particularly important because this pathology colocalizes with markers of the translational stress response, such as stress granule proteins HNRNPA2B1, TIA1, EIF3n and PABP, and also neuronal death (as shown by cleaved caspase 3 and loss of NeuN positive cells) [23, 25, 36, 51-53]. Thus, neurodegeneration appears to be potentiated in the APP^NL-G-F^/MAPT^P301S^ model.

Increasing evidence points to a key role for inflammation in AD. Many AD-linked genes appear to enhance disease risk by affecting the biology of microglia. For instance, TREM2 is one of the strongest risk factors for AD, and evidence suggests that TREM2 acts to recognize Aβ pathology and direct microglial responses toward the pathology [54, 55]. The strong impact of Aβ on inflammation is also evident in the APP^NL-G-F^/MAPT^P301S^ model. Inflammatory cells are readily evident in the area around Aβ plaques (Fig. 3A). These microglia were more abundant and showed greater ramifications than in the P301S MAPT model. However, it is important to note that microglia do respond to MAPT pathology; Iba1 reactivity was increased in the P301S MAPT model, but not to the same level as observed in the APP^NL-G-F^ or APP^NL-G-F^/MAPT^P301S^ model. These results are consistent with published work indicating that Aβ pathology elicits a phenotype termed “Disease Associated Microglia” while MAPT pathology elicits microglial responses exhibiting weaker cytokine production [56].

The increases observed for m^6^A parallel those reported by our group previously [23]. We observed that increases in m6A were observed most prominently in neurons, and that levels of m^6^A correlated with levels of MAPT pathology. This finding is consistent with the observation that MAPT pathology co-localizes with the RNA binding protein, HNRNPA2B1, which functions as an indirect m6A reader. The results presented above in Figure 5 directly compare m^6^A levels in the context of MAPT pathology, Aβ pathology and combined MAPT and Aβ pathology. The results are clear in that m^6^A levels follow MAPT pathology. A previous manuscript examining m^6^A in an AD model utilizing over-expression of only APP, which is a model that exhibits Aβ accumulation without corresponding MAPT pathology. The data presented above suggest that the absence of MAPT pathology in this model produced a corresponding absence of m^6^A accumulation [44]. Our studies also provide insight into potential mechanisms regulating m^6^A in disease. The increases in m^6^A observed correlate with MAPT pathology are show a corresponding association with increased levels of METTL3 and reduced levels of ALKBH5, which are the enzymes that respectively add and remove m^6^A from mRNA. More writing combined with reduced erasure leads to increased levels of m6A.

Although changes in m^6^A were most evident in neurons, the results in Figure 5A show that some astrocytes and microglia also exhibited strong increases in m^6^A in the APP^NL-G-F^/MAPT^P301S^ model. Such results are consistent with emerging studies showing that m^6^A regulates inflammation, macrophages and also astrocytosis [57-61]. Such findings raise the possibility that m^6^A modulation might also be applied towards regulation of inflammation in AD; indeed a recent study observed that conditional knockout of METTL3 in microglia attenuated inflammation and Aβ accumulation in a mouse model based on Aβ injection [62].

## Conclusion

The field is increasingly moving towards use of KI models. The APP^NL-G-F^/MAPT^P301S^ model described in this manuscript takes advantage of the APP^NL-G-F^ KI line, but utilizes the human MAPT^P301S^ line in order to achieve robust MAPT pathology. Our results show that this model provides an appealing alternative to the double APP^NL-G-F^ x MAPT KI model, which develop both MAPT pathologies very slowly [21]. The current APP^NL-G-F^/MAPT^P301S^ model thus provides a useful compromise. This model exhibits strong Aβ accumulation and strong inflammation, while avoiding artifacts associated with APP or presenilin over-expression. The APP^NL-G-F^/MAPT^P301S^ model also possesses the benefits arising from strong expression of human MAPT with the resulting rapid development of robust MAPT pathology, strong m^6^A accumulation and, importantly, significant neurodegeneration.

## Abbreviations

Aβ: β-Amyloid
AD: Alzheimer disease ALKBH5: AlkB Homolog 5 APP: Amyloid precursor protein
ELISA: Enzyme-linked immunosorbent assay FTO: Fat mass and obesity associated
GFAP: Glial fibrillary acidic protein
HNRNPA2B1: Heterogeneous Nuclear Ribonucleoprotein A2/B1 Iba-1: Ionized calcium binding adaptor molecule 1
m^6^A: N^⁶^-Methyladenosine
MAPT, Tau: microtubule associated protein tau Mettl3: Methyltransferase-like 3
qPCR: Quantitative polymerase chain reaction
YTHDF1: YTH N^6^-Methyladenosine RNA Binding Protein 1 YTHDF2: YTH N^6^-Methyladenosine RNA Binding Protein 2

## Availability of data and materials

Generated datasets used for analyses in this study are available from the corresponding author upon reasonable request.

## Acknowledgements

We would like to thank the following funding agencies for their support: BW: NIH (AG050471,

R01AG080810, AG056318, AG064932, AG061706, UO1AG072577) and the BrightFocus

Foundation. SR: JSPS Kakenhi 20KK0338.

## Author contributions

Conceptualization: B.W. and L.J.;

Methodology: L.J., R.R., W.X., T.S., and T.C.S., S.R., J.S., and P.C.D.;

Investigation and Visualization: L.J., R.R., M.W., L.Z., C.J.W., A.K., M.J, J.S. and S.A.D.; Data Analysis and Writing: L.J. and B.W.;

Supervision and Funding Acquisition: B.W.

## Declaration of Interests

B.W. is co-founder and Chief Scientific Officer for Aquinnah Pharmaceuticals Inc. All other authors declare that they have no competing interests to disclose.

